# Structural variants identified by Oxford Nanopore PromethION sequencing of the human genome

**DOI:** 10.1101/434118

**Authors:** De Coster Wouter, De Roeck Arne, De Pooter Tim, D’Hert Svenn, De Rijk Peter, Strazisar Mojca, Kristel Sleegers, Van Broeckhoven Christine

## Abstract

We sequenced the Yoruban NA19240 genome on the long read sequencing platform Oxford Nanopore PromethION for benchmarking and evaluation of recently published aligners and structural variant calling tools. In this work, we determined the precision and recall, present high confidence and high sensitivity call sets of variants and discuss optimal parameters. The aligner Minimap2 and structural variant caller Sniffles are both the most accurate and the most computationally efficient tools in our study. We describe our scalable workflow for identification, annotation, and characterization of tens of thousands of structural variants from long read genome sequencing of an individual or population. By discussing the results of this genome we provide an approximation of what can be expected in future long read sequencing studies aiming for structural variant identification.

## Introduction

Structural variants (SVs) are defined as regions of DNA larger than 50 bp showing a change in copy number or location in the genome, including copy number variants (CNVs, deletions and duplications), insertions, inversions, translocations, mobile element insertions, expansion of repetitive sequences and complex combinations of the aforementioned (Escaramís et al. 2015; Sudmant et al. 2015). SVs account for a higher number of variable nucleotides between human genomes, even though single nucleotide variants (SNVs) are far more numerous (Conrad et al. 2010). However, the majority of SVs are poorly assayed using currently dominant short read sequencing technologies but can be detected using long read sequencing technologies from Pacific Biosciences and Oxford Nanopore Technologies (ONT) (Chaisson et al. 2015, 2018). Long read sequencing technologies have a lower raw accuracy of approximately 85% but have the advantage of a better mappability in repetitive regions, further extending the part of the genome in which variation can be called reliably (Li and Freudenberg 2014).

Sequencing DNA fragments using a protein nanopore is a relatively old concept, which got commercialized by ONT with the release of the MinION sequencer five years ago (Deamer et al. 2016). A MinION flow cell has 512 sensors collecting measurements from 2048 pores. This device’s minimal initial investment, long reads and rapid results has enabled many applications for smaller genomes (Loman et al. 2015; Quick et al. 2015; Jansen et al. 2017; Bainomugisa et al. 2018;Miller et al. 2018). Recent runs routinely reach 8 Gbase and currently up to 30 Gbase, with a big in-field variability and incremental improvements over the years (Schalamun et al. 2018). Applications for human genomics could only be achieved by combining multiple flow cells, which is cumbersome and costly. Early adopters investigated structural variants in two genomes from patients with a congenital disorder due to chromothripsis by combining data from 135 flow cells (Cretu Stancu et al. 2017), and a consortium of MinION users sequenced and released data from the human reference sample NA12878 generated on 39 flow cells reaching 91.2 Gbase or ~30x coverage (Jain et al. 2018).

Routine human genome sequencing applications have become possible on the recently commercially available PromethION sequencer. A PromethION flow cell has 3000 sensors and 12000 pores, which generates on average 70 Gbase of data in our hands <u>(De Roeck et al. 2018)</u>, allowing for the sequencing of a 20x covered human genome per flow cell. At the moment up to 24 flow cells can be run simultaneously on the machine, with a future upgrade to 48 planned. Here, we present the characteristics of such PromethION runs, and a bioinformatic workflow for identification and characterization of structural variants. Finally, we provide a detailed description of the Yoruba NA19240 reference sample compared with publicly available variant data and discuss implications for future structural variant detection projects from long read sequencing.

## Results

### Human genome sequencing on PromethION

We generated a 59x median genome coverage of NA19240 on PromethION by combining data from five flow cells, which we compared with MinION data from the same sample. The run metrics and ENA accession IDs are summarized in Table 1 and Supplementary Figure S2. The longest read we obtained was 177 kb on PromethION and 219 kb on MinION. Overall, our results are suggestive of an inverse relationship between yield and read lengths, as higher yields were obtained for libraries which included shearing to 20 kb fragments.

**Table 1:** library characteristics and accession ids.

| Library identifier | Yield [Gbase] | Number of reads [millions] | Median read length [kb] | N50 [kb] | ENA accession id |
| --- | --- | --- | --- | --- | --- |
| P1-N | 48.3 | 3.5 | 12.4 | 20.0 | ERR2631600 |
| P2-S | 58.4 | 4.9 | 11.9 | 14.1 | ERR2631601 |
| P3-N | 22.2 | 1.4 | 13.7 | 29.1 | ERR2631602 |
| P4-N | 19.6 | 1.2 | 13.8 | 28.4 | ERR2631603 |
| P5-S | 59.1 | 6.2 | 9.8 | 11.8 | ERR2631604 |
| M1-N | 4.7 | 0.4 | 11.6 | 19.6 | ERR2683660 |
| M2-N | 7.7 | 0.3 | 20.4 | 32.8 | ERR2683661 |

### Comparing MinION and PromethION

The obtained read lengths were similar between matched library preps sequenced on MinION and PromethION (respectively P1-N and M1-N, P3-N and M2-N) (Figure 1A, B). We observed a higher average quality score and corresponding percent identity after alignment to the human reference genome GRCh38 for the MinION data (p<0.0001, Mann-Whitney U test) (Figure 1C, D).

**Figure 1:**
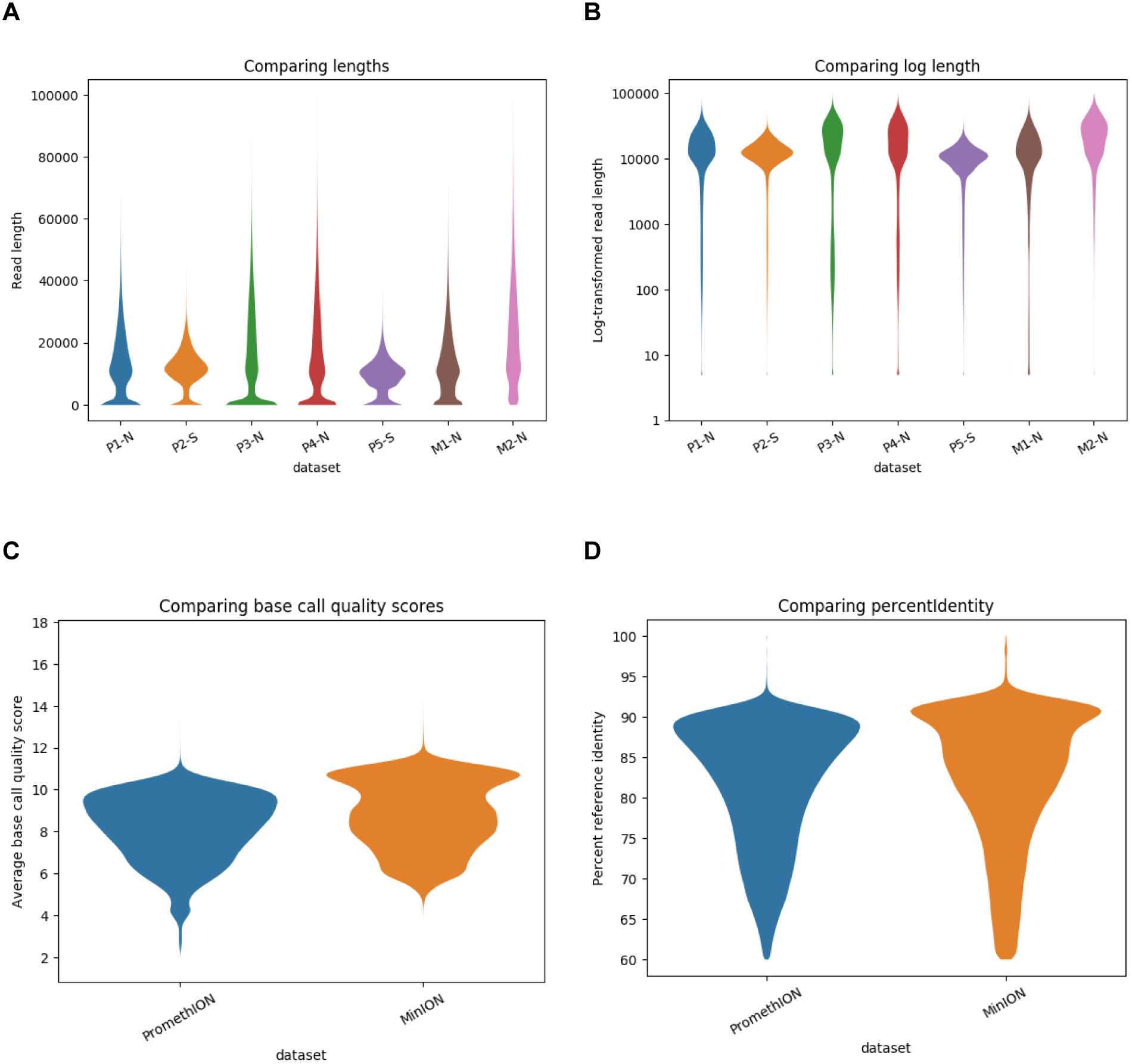
Comparison of PromethION and MinION libraries. A: read lengths; B: log-transformed read lengths; C: per read average base call quality scores; D: percent identity after alignment to the reference genome P: PromethION; M: MinION; N: non-sheared/native; S: sheared before library preparation Plots were made using NanoPack (De Coster et al. 2018).

### Comparing aligners

The characteristics of the alignments using ngmlr (Sedlazeck et al. 2018), LAST (Kiełbasa et al. 2011) and two parameter settings of minimap2 (H.Li 2018) are shown in Table 2 and Figure 2 and S3. LAST generates many split alignments leading to more, shorter aligned reads. ngmlr favors aligning short exact fragments for a subset of the reads (Figure 2). Percent identity comparisons are roughly equivalent, with a slightly lower median for LAST. Minimap2 is the fastest and LAST by far the slowest of the three aligners. Median alignment coverage is approximately equal, with the highest coverage by ngmlr and the lowest by LAST. 90% of the genome is covered at 40x (Figure S3).

**Table 2:** metrics of aligners.

| Aligner | Gigabases aligned | Median length | Median coverage | Runtime* [seconds] |
| --- | --- | --- | --- | --- |
| ngmlr | 178.3 | 11573 | 59 | 306 |
| LAST | 169.2 | 1240 | 56 | 4470 |
| minimap2 | 187.1 | 11165 | 58 | 46 |
| minimap2-last-like | 179.5 | 7506 | 57 | 59 |

\* Average of three measurements on 10 000 reads, aligned using 12 threads.

**Figure 2:**
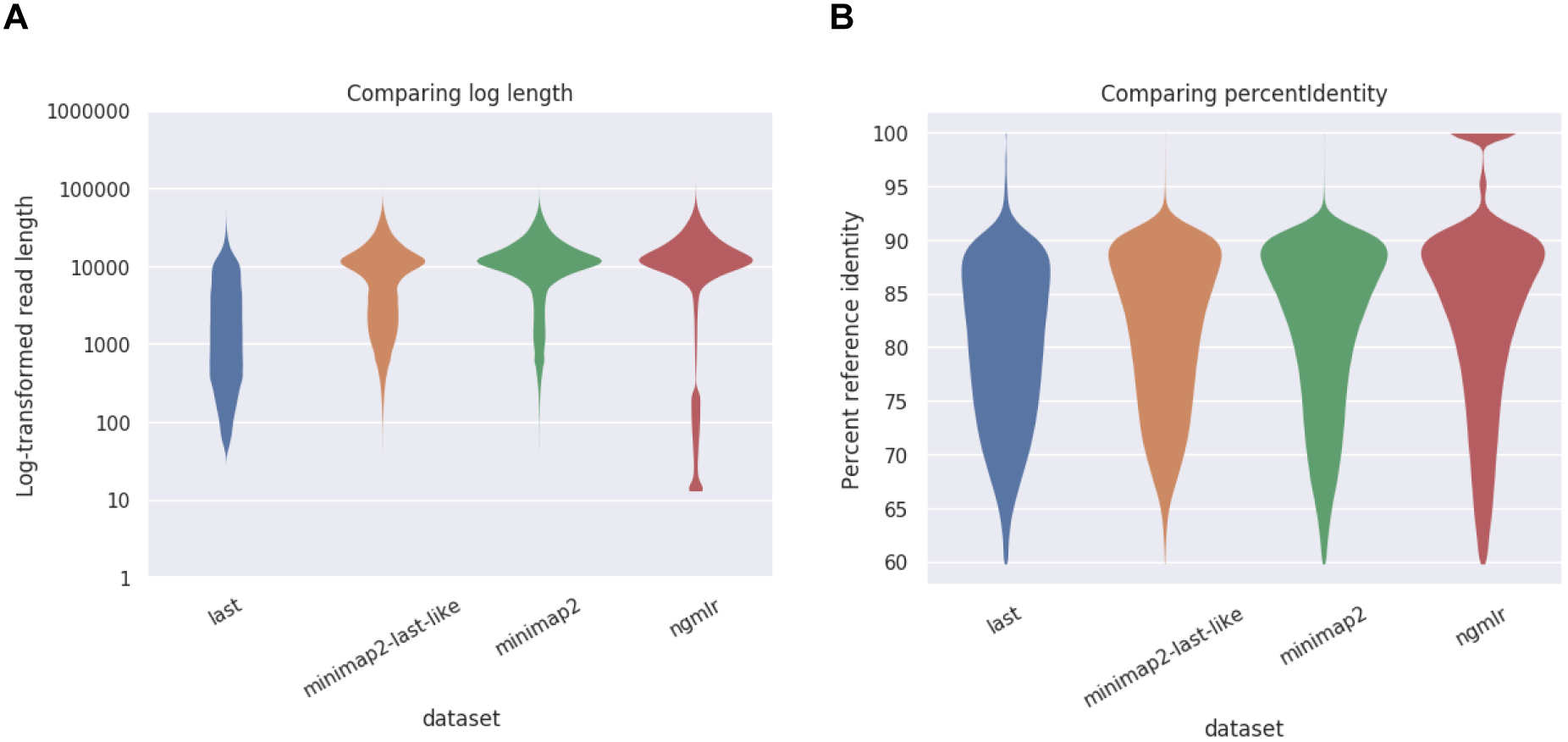
Comparison of aligners. A: log-transformed aligned read lengths; B: read percent identity compared to the reference genome. Plots were made using NanoPack (De Coster et al. 2018).

### Structural variant calling

Structural variants were called using Sniffles (Sedlazeck et al. 2018) and NanoSV (Cretu Stancu et al. 2017), and inversions additionally with npInv (Shao et al. 2018). The number of variants identified and the runtime per dataset are summarized in Table 3. A detailed overview of the number of variants split by variant type can be found in Supplementary Table 2. The truth set of SVs from NA19240 contains 29436 variants, of which 10607 deletions, 16337 insertions and 2503 variants of other types, such as repeat expansions. NanoSV consistently identifies more variants, with a strikingly high number after LAST alignment. LAST alignment turned out to be incompatible with calling variants using Sniffles and npInv.

Sniffles is by far the fastest of the evaluated SV callers. In contrast, we stopped NanoSV after one month when using the full dataset, which also required > 100 Gbytes of RAM per sample, presumably since it is not yet adapted to this volume of data. To circumvent this issue in our workflow alignments are split by chromosomes and processed in parallel.

**Table 3:** metrics of SV callers.

| Aligner | SV caller | Number of variants | Relative run time* |
| --- | --- | --- | --- |
| minimap2 | Sniffles | 28305 | 1.18 |
|  | NanoSV | 64761 | 94.07 |
|  | nplnv | 167 | 2.04 |
| minimap2-last-like | Sniffles | 15821 | 1 |
|  | NanoSV | 38182 | 80.36 |
|  | nplnv | 419 | 2.71 |
| ngmlr | Sniffles | 29159 | 1.02 |
|  | NanoSV | 47431 | 60.25 |
|  | nplnv | 461 | 2.27 |
| LAST | NanoSV | 123729 | 257.98 |

\*Relatively to the fastest, tested using 12 threads, on a subset of the data using an average of three measurements

### Precision-recall and combined SV sets

We evaluated the precision and recall of identified SVs for combinations of the described aligners and SV callers (Figure 3A and Supplementary Table 3). All combinations perform approximately in the same range, with the exception of Sniffles after minimap2 alignment with last-like parameters and NanoSV after LAST alignment, which respectively showed a mediocre recall and precision. Of note, Sniffles performed at a higher precision with similar accuracy after minimap2 alignment compared with ngmlr alignment. For this reason, further parameter evaluations will be performed using the combination of Sniffles after minimap2 alignment. For the ngmlr alignment NanoSV performed better than Sniffles. There is also a considerable number of SV calls overlapping between Sniffles and NanoSV after alignment using minimap2, which are not part of the truth set (Supplementary Figure S4). A similar analysis using manta (Chen et al. 2016) and lumpy (Layer et al. 2014) after bwa mem alignment (H.Li 2013) of Illumina reads was performed. Manta performed the best, with a precision of 55% and recall of 28% with 15122 identified SVs, while lumpy only reached 18% precision and 4% recall with 6100 identified SVs.

**Figure 3:**
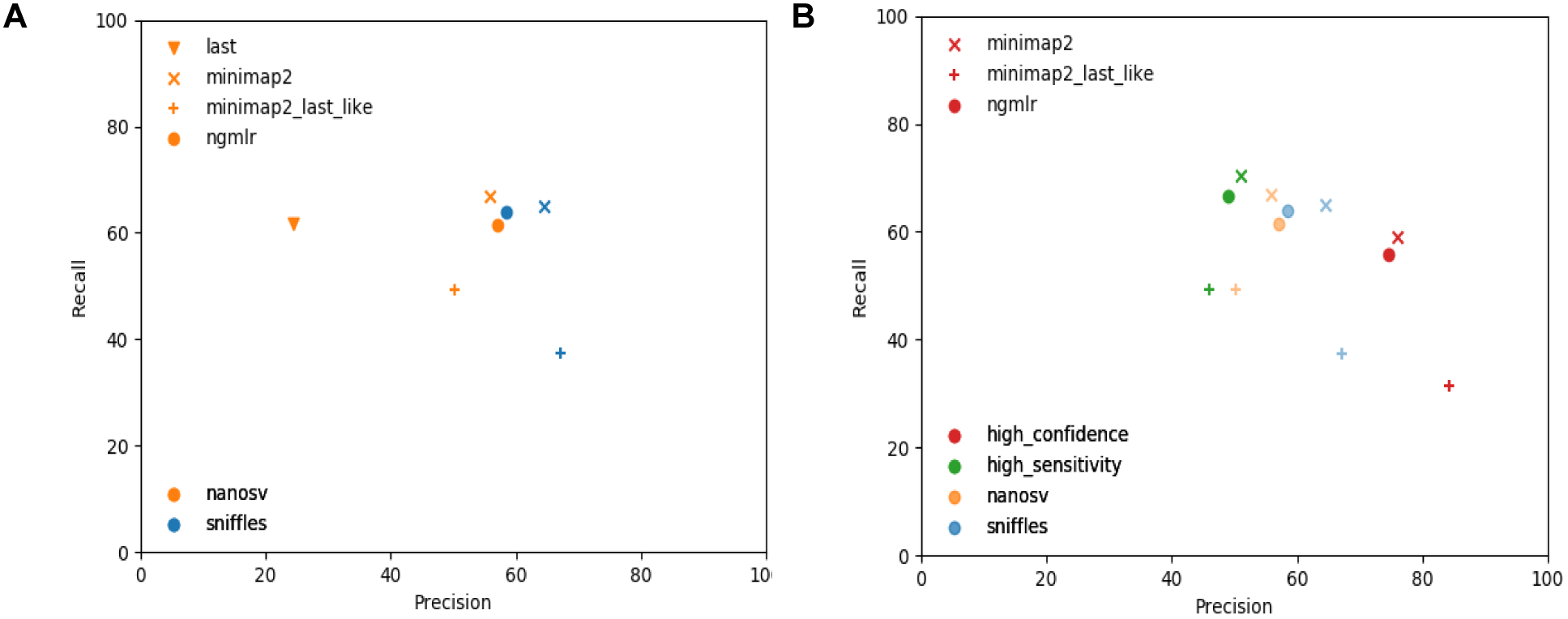
Precision-recall comparison. A: Precision and recall of individual callsets; B: including combined call sets Aligners are tagged with symbols, variant callers and combined sets are tagged with colors.

We separately evaluated inversions called by Sniffles, NanoSV and the specifically tailored tool npInv. As breakpoints of inversions are typically associated with long repetitive sequences, leading to inaccurate alignments, we allowed a larger distance between pairs of breakpoints up to 5kb to be considered concordant. The accuracy is generally poor and the best results were obtained using npInv, with precision between 17.8% and 33.8% and recall between 19.9% and 25.4% (Supplementary Table 3).

High-confidence and high sensitivity variant sets based on respectively the intersection and union of Sniffles and NanoSV were generated for the aligners used in this comparison. Their precision and recall compared with the truth set are shown in Figure 3B. No combined call sets were made for LAST alignment since no variants can be called using Sniffles. Alignments performed with minimap2 consistently outperform those with ngmlr in both sensitivity and confidence. A remarkable outlier was again alignment with last-like parameters for minimap2, which showed a high precision but low recall.

We also evaluated the accuracy of the zygosity determination of Sniffles and NanoSV after minimap2 alignment (Table 4), showing that these are often inaccurate. For NanoSV these results did not change notably when limiting to those SVs for which divergent sequencing depth was available to support the variant type classification (data not shown). When evaluating precision-recall separately for either insertions or deletions we observed no remarkable difference for NanoSV, but it turned out that Sniffles has a lower precision and higher recall for a loss of sequence and conversely for gains of sequence a higher precision but lower recall (Supplementary Table 3). As such insertions turned out to be more often correct than deletions, while proportionally more deletions from the truth set were identified.

**Table 4:** Accuracy of zygosity.

|  |  | Sniffles |  |  | NanoSV |  |  |
| --- | --- | --- | --- | --- | --- | --- | --- |
|  |  | nocall | het | hom | nocall | het | hom |
| Truth | nocall | 0 | 3295 | 5828 | 236 | 11972 | 1433 |
|  | het | 4535 | 4151 | 5837 | 4064 | 9807 | 652 |
|  | hom | 4544 | 321 | 2947 | 4894 | 2464 | 3454 |

A parameter of Sniffles determines the minimum number of reads supporting an SV before it gets reported, with 10 as the default. Testing multiple values for this demonstrated a clear trade-off between precision and recall, as shown in Figure 4A. When allowing less support for a candidate SV the recall was the highest, but precision was low and vice versa. An appropriate middle ground appeared to be around ¼ or ⅓ of the median genome coverage to maximize precision and recall.

By randomly down-sampling the alignment from minimap2 to various fractions of the original dataset we evaluated the influence of the median genome coverage on the precision and recall (Figure 4B). Sniffles was used with default parameters (minimal_support = 10). We concluded that increasing the coverage above 40x did not substantially increase the recall. The reduction in precision above that value originated in a sub-optimal selection of the minimal support parameter as described above.

**Figure 4:**
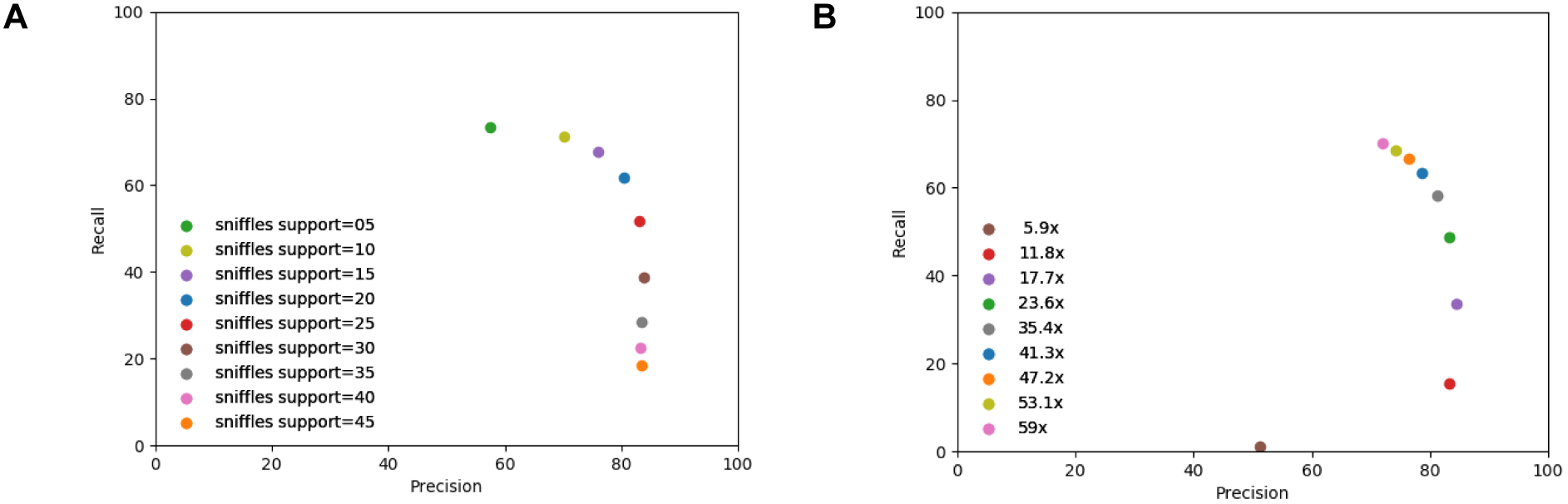
precision and recall with parameter variation. A: Specifying minimally supporting reads; B: influence of the median genome coverage after down-sampling to various fractions. Both sets use Sniffles SV calling and minimap2 alignment.

We also evaluated the length of SVs and their accuracy (Figure 5 and Supplementary Figure S5). A substantial proportion of variants <100 bp, the largest group, turned out to be either missed or false positive by both Sniffles and NanoSV. Most of the variants correctly identified by Manta were <300 bp and compared to the long read SV callers more variants were missed in each length category (Supplementary Figure S6).

**Figure 5:**
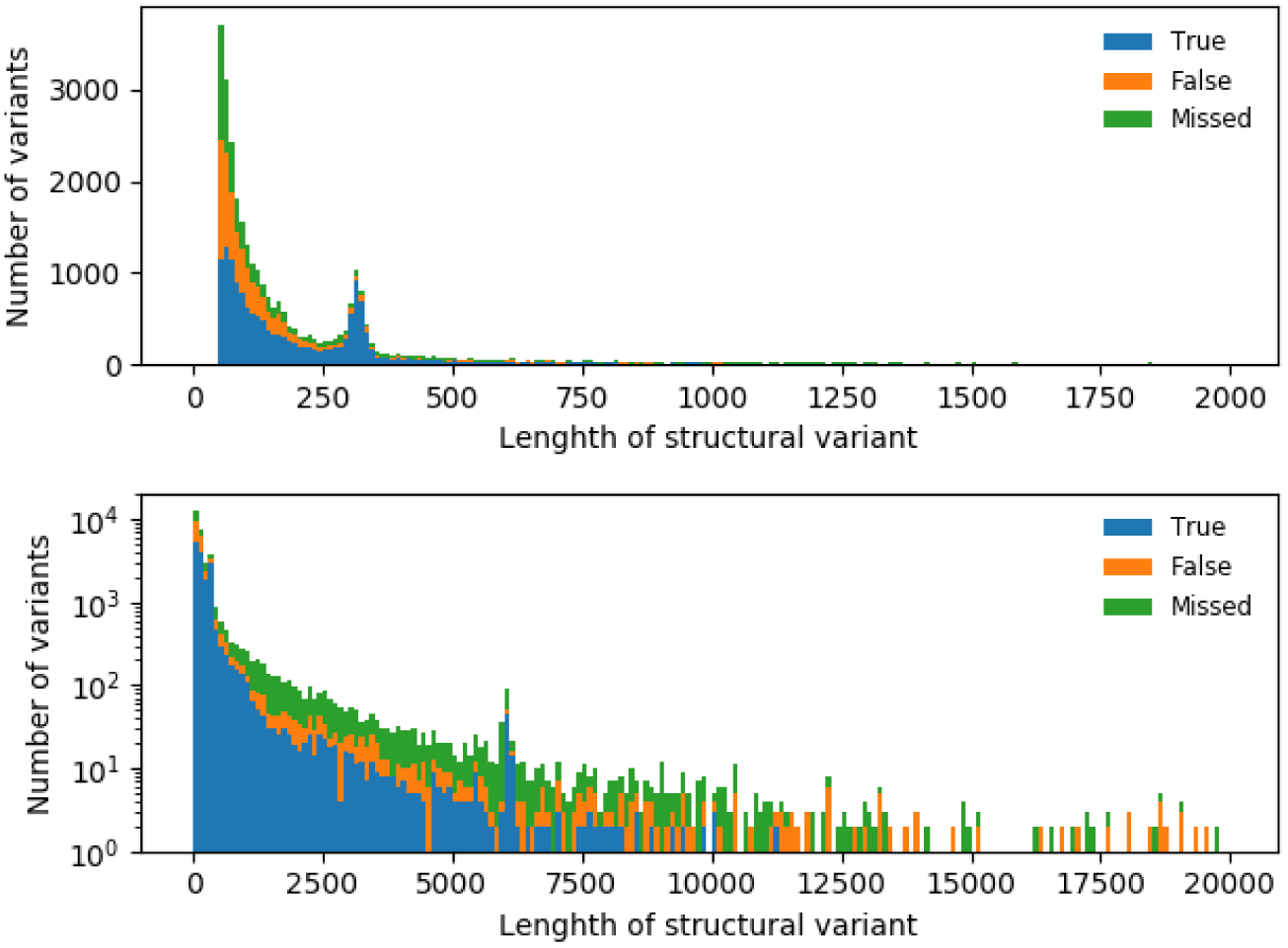
SV validation status per length. SVs identified using Sniffles after minimap2 alignment, compared to truth set

### Description of detected variants

Sniffles after minimap2 alignment detected 28305 SVs. Of those 11822 overlapped with genes, of which 695 were in coding sequences. 3780 variants overlap with segmental duplications. The profile of lengths of SVs in the truth set and after SV calling with Sniffles and NanoSV (Figure 6 and Supplementary Figure S5) were comparable, showing a peak around 300 bp corresponding to SVs involving Alu elements and around 6 kb corresponding to L1 elements, an observation also reported in other studies (Huddleston et al. 2017; Cretu Stancu et al. 2017).

**Figure 6:**
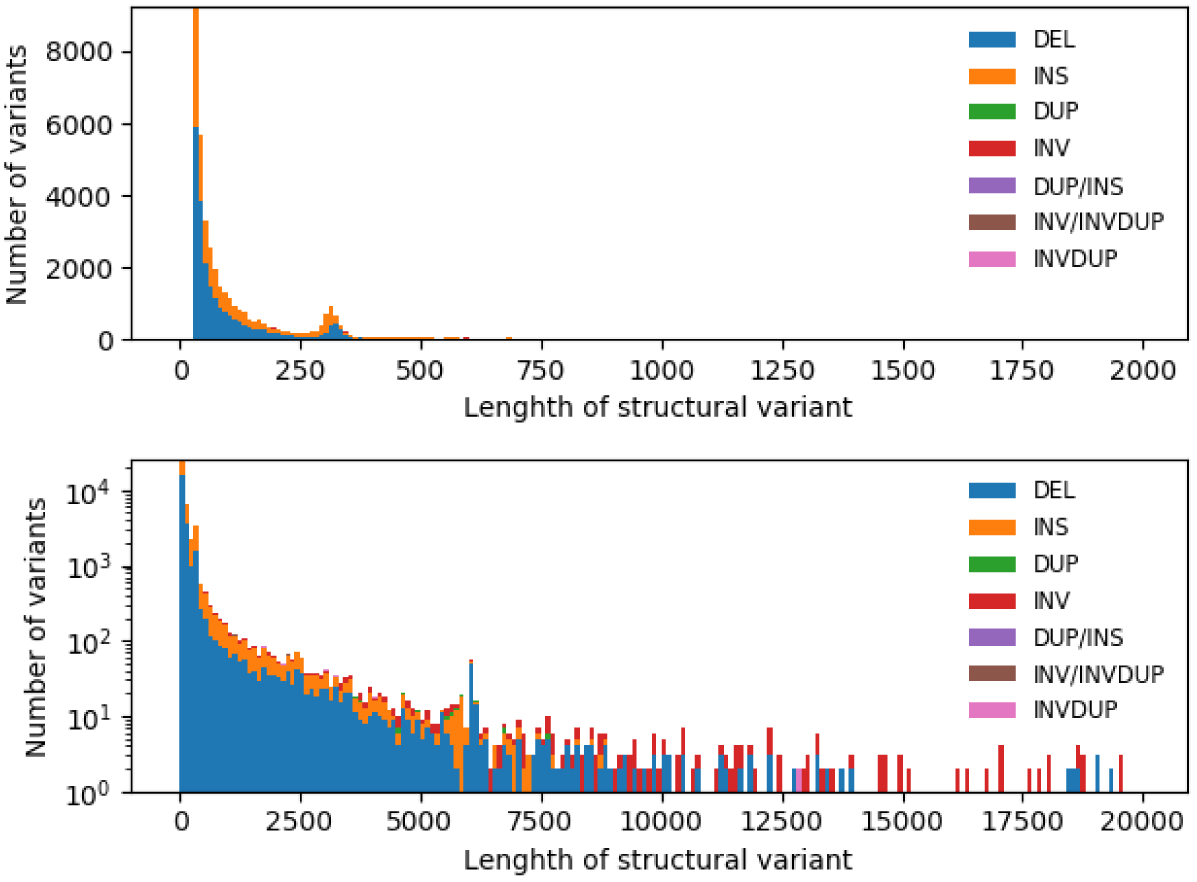
length profile of SV calls. SV calls made by Sniffles after minimap2 alignment. The upper panel has SVs up to 2 kb, the lower up to 20 kb with a log-transformed number of variants.

## Discussion

For benchmarking and characterization we sequenced a well-characterized reference genome on the Oxford Nanopore PromethION. We observed a substantial variability in the yield of the PromethION which can partially be attributed to using sheared or unsheared DNA and differences in concentration of the final library. Further optimizations of the library preparation will point to the optimal balance between yield and read length. While much longer reads have been reported by other users on MinION (Payne et al. 2018), this was not our primary aim. Based on the higher quality scores and percent identity it seems that the MinION system is currently better understood, yielding slightly more accurate results. However, we assume similar improvements can be made to the PromethION base calling models when more data becomes available and the system has matured to the level of the MinION. In fact, an updated version of the basecaller is available, modestly improving the nucleotide-level accuracy. However, we do not anticipate a large impact on SV detection. Further improvements in base call quality might increase the accuracy of breakpoint detection.

In this work, we evaluated the precision (positive predictive value) and recall (sensitivity) for multiple long read aligners and SV callers. While the truth set in our comparison was based on multiple integrated technologies (Chaisson et al. 2018), we cannot exclude that any variants were missed. However, with the combination of short read, long reads, linked reads and strand-seq both shorter and longer SVs, and more challenging inversions have been characterized. For this application, we anticipated this set was sufficiently complete. Follow-up of high-confidence, but presumably false positive variants using orthogonal methods could be valuable.

Ngmlr uses a convex gap cost allowing alignment across SVs, while simultaneously accounting for small indels, the dominant errors in long read sequencing. Alignments are split in the case of larger SVs. This aligner is co-developed with the SV caller Sniffles (Sedlazeck et al. 2018). Minimap2 is an accurate and ultra-fast aligner for reads from (paired) short and long DNA sequencing, long RNA sequencing or assembled contigs and genomes. SVs between reference and individual are taken into account using a concave gap cost (Li 2018). LAST was not developed recently but was suggested to be the most accurate for e.g. NanoSV (Cretu Stancu et al. 2017). LAST is developed for sensitive comparison of long sequences and is based on BLAST but reduces alignment time by eliminating seeds in repetitive elements (Kiełbasa et al. 2011). However, we demonstrated that it is clearly not optimal for this volume of data and takes too long to be used on a larger scale.

Sniffles calls indels, duplications, inversions, translocations, and nested SVs and takes read support, SV size and breakpoint consistency into account (Sedlazeck et al. 2018). Unexpectedly, we observed that Sniffles, while developed for ngmlr alignment, performed better after minimap2 alignment, yielding a higher precision at a similar recall compared to ngmlr. Even higher precision was obtained using the ’last-like’ parameter settings of minimap2, but this came at a considerable trade-off of lower recall and is therefore not recommended. NanoSV uses clustering of split and gapped alignments for identification of breakpoint junctions, (Cretu Stancu et al. 2017). While Sniffles is very fast, NanoSV requires further software optimizations to handle these large volumes of data and limit runtime and memory usage. We had to execute NanoSV per chromosome in parallel to keep the runtime reasonable, with the limitation that interchromosomal variants cannot be detected. However, for our application of germline structural variants in a healthy individual, we expected these to be less relevant. While LAST is the recommended aligner by the authors of NanoSV the number of variants identified was excessive, with a high number of false positives. NanoSV, too, obtained the best results after minimap2 alignment. In our comparison with SVs called from short read sequencing data using Manta and Lumpy a clear advantage for long reads was demonstrated, with substantially higher recall values. However, our evaluation of the accuracy of the zygosity of the identified SVs showed that these are highly unreliable, owing to the complexity of a diploid genome. We also observed that both SV callers performed poorly in the largest group of variants, those with a length below 100 bp. Thousands of variants in this group were either missed (not identified) or false positively called as SV. Improvements in this size range, or more specialized SV callers, are definitely welcome here.

Inversions are generally challenging or impossible to identify, especially using traditional methods such as comparative genome hybridization, PCR-based approaches or short read sequencing, because those variants are copy-neutral with breakpoints often in (long) highly repetitive sequences. For a comprehensive detection Strand-Seq is currently the only applicable protocol (Chaisson et al. 2018). Long read sequencing could offer an advantage, as these might provide more accurate alignments in repetitive sequences. However, we observe here a generally low precision and recall for the callers identified, with precision and recall between around 15 and 30%, with much lower precision for NanoSV. We hypothesize that even longer read lengths might be beneficial here, together with algorithmic improvements.

Due to its speed, we could evaluate relevant parameters for Sniffles, and concluded that adding more than 40x coverage did contribute little to the identification of novel variants. Presumably longer reads might reveal more hidden variation in highly repetitive sequences. We suggest using a minimal supporting number of reads of one-fourth to one-third of the median genome coverage to optimize precision and recall. Ultimately, setting stringency filters is a trade-off between sensitivity and specificity, for which the applications at hand determine if it’s appropriate to tolerate false positives, or rather accept that some genuine calls can be missed. It is worth noting that Sniffles can be used in a two-pass mode, in which variants identified in the first stage can be used to force genotyping in a second stage. As such this shifts the burden of ’discovery’ of SVs to ’genotyping’ known SVs, potentially increasing the sensitivity in larger cohorts and at lower coverage.

After the identification of SVs, we also annotated these with information about overlapping genes, segmental duplications and known variants in DGV. Obviously overlapping genes are relevant to judge the potential pathogenicity of the identified variants, while the impact of SVs in non-coding regions is currently less well understood. The annotation of SVs localized in a segmental duplication plays a double role, as these regions are both known to be a hotspot for SV formation, but simultaneously are troublesome for alignments and as such can give rise to false positive variant calls (Stankiewicz et al. 2003; Sharp et al. 2005; Bailey and Eichler 2006).

Here we provided an estimate of what can be expected in future long read whole genome sequencing data. While SVs can contribute to disease, it was clear that, just as with the better understood SNVs, the majority will be mostly harmless. To distinguish pathogenic SVs from polymorphisms we will need comprehensive catalogs across multiple populations.

## Methods

### Sample preparation

The lymphoblastic cell line (LCL) GM19240 was ordered from the Coriell Cell Repository and cultured as specified by the Coriell Institute for Medical Research (Camden, NJ USA). DNA from GM19240 (NA19240) was extracted using both manual QIAmp DNA Blood mini kit (Qiagen, USA) and a robotic extraction platform (Magtration system 8LX, PSS, Japan), as specified by the suppliers. As specified in the QIAamp protocol RNase A treatment was performed during the extraction, while robotically extracted DNA was treated additionally with RNase A (RNase A, 10mg/ml, using 1μl RNase A per 100μl template, Thermo Fisher Scientific, USA) to remove RNA.

Extractions of 5 million LCL cells, resuspended in 200 μl of PBS, as well as extraction with 8LX (PSS, JP), yielded between 15 to 17 μg of gDNA per extraction, with average A260/280 of 1.86, A260/230 of 2.50 and average gDNA size between 38 and 41 kb. Information on the five aliquots used for library preparation is supplied in the Supplementary Table 1. The fifth aliquot (P5) was a pool from DNA of automated PSS and manual QIAmp extraction.

As we wanted to evaluate the efficiency of different library preparations, two out of five aliquots were sheared using Megaruptor (Diagenode, BE) to an average size of 20 kb and three aliquots were non-fragmented. All aliquots were purified and size selected using a high pass protocol and the S1 external marker on the BluePippin (on 0.75% agarose gel, loading 5 μg sample per lane) (Sage Science, MA, USA). The size selection cutoff differed between fragmented and unfragmented samples (Supplementary Table 1). The average recovery of the size-selected DNA aliquot was between 40-70% of the initial input. After size selection, all aliquots were purified using AMPure XP beads (Beckman Coulter, USA) using ratio 1:1 (v:v) with DNA mass recoveries between 88-99%. All fragment analyses were performed on Fragment Analyzer with DNF-464 High Sensitivity Large fragment 50 kb kit, as specified by the manufacturer (Advanced Analytical, Agilent, USA).

### Library preparation

The recommended protocol for library preparation on PromethION was followed with minor adaptations. In short, potential nicks in DNA and DNA ends were repaired in a combined step using NEBNext FFPE DNA Repair Mix and NEBNext Ultra II ER/dAT Module (New England Biolabs, USA) followed by AMPure bead purification and ligation of sequencing adaptors onto prepared ends. Four libraries were constructed using 1D DNA Ligation Sequencing kit SQK-LSK109 following the PromethION protocol (GDLE_9056_v109_rev E_02Feb2018) and one (P4) was made using Ligation Sequencing kit SQK-LSK108 following the SQK-LSK108- PromethION protocol (GDLE_9002_V108_revO_28Mar2018) since LSK109 consumables were depleted at that time. The main differences between SQK-LSK109 and SQK-LSK108 protocols are increased ligation efficiency, a different clean-up step, the combined FFPE repair and end-repair. These modifications, making sequencing of long reads more efficient, were used for both protocols.

Additionally to consumables supplied with the sequencing kit, several steps were performed using NEB enzymes (NEBNext FFPE DNA Repair Mix, NEBNext Ultra II ER/dAT Module and NEBNext Quick Ligation Module, all New England Biolabs, USA) as recommended in 1D genomic ligation protocols (SQK-LSK 109 and SQK-LSK108). Overall, ONT protocols were followed, with slight increases in incubation times during DNA template end-preparation, purification, and final elution. The final mass loaded on the flow cells was determined based on the molarity, depended on average fragment size and was chosen based on our prior experience and communication with specialists at Oxford Nanopore Technologies.

Two aliquots of the unfragmented NA19240 were used for library preparation and sequencing on MinION using R9.4.1 flow cells as a quality control for library preparation (Oxford Nanopore Technologies, UK) (Supplementary Table 1). The MinION libraries M1-N and M2-N were prepared identically to PromethION libraries ’P1-N’ and ’P3-N’, respectively.

### Data processing

Base calling of the raw reads from MinION and PromethION was performed using the Oxford Nanopore basecaller Guppy (v1.4.0) on the PromethION compute device. Run metrics were calculated, summarized and compared to each other using NanoPack (De Coster et al. 2018). Reads were aligned to GRCh38 from NCBI, without alternative contigs and including a decoy chromosome for the Epstein–Barr virus (H.Li 2017)^1^. Reads were aligned with ngmlr (v0.2.6) (Sedlazeck et al. 2018), LAST (v876) (Kiełbasa et al. 2011) and two parameter settings of minimap2 (v2.11-r797), of which one was supposed to mimic alignments by LAST (H.Li 2018)

(see Supplementary data for commands and parameters). The substitution matrix for alignment with LAST was determined using LAST-TRAIN with a subset of 5000 reads (Hamada et al. 2017). Coverage was assessed using mosdepth (Pedersen and Quinlan 2018). Processes were parallelized using gnu parallel (Tange 2011).

### Comparison of structural variant calls

Structural variant calling was performed using Sniffles (v1.0.8) (Sedlazeck et al. 2018) and NanoSV (v1.2.0) (Cretu Stancu et al. 2017) with default parameters. Inversions were called with the specific tool npInv (Shao et al. 2018). Alignment with LAST turned out to be incompatible with Sniffles and npInv. We were unsuccessful at using Picky (Gong et al. 2018) and reported several issues to the authors, which remained unanswered and unresolved. For reference, we also evaluated the short read SV callers Manta (Chen et al. 2016) and Lumpy (Layer et al. 2014) after bwa mem alignment (H.Li 2013) from Illumina data generated by Chaisson et al (Chaisson et al. 2018).

As a gold standard truth set of SVs in NA19240, we used haplotype-resolved SVs which were identified by combining PacBio long read sequencing, Bionano Genomics optical mapping, Strand-Seq, 10x genomics, Illumina synthetic long reads, Hi-C and Illumina sequencing libraries (Chaisson et al. 2018). This set of variants will be called the "truth set" from here on. It is worthy of note that both Sniffles and NanoSV report SVs from at least 30 bp, while the formal definition and the truth set use 50 bp as the lower limit. Therefore all accuracy calculations are performed for variants ≥ 50 bp. Obtained variant call sets were combined SURVIVOR (v1.0.2) (Jeffares et al. 2017) and precision-recall metrics were calculated after parsing with cyvcf2 (Pedersen and Quinlan 2017) and plotted using matplotlib (Hunter 2007). For combining SVs a distance of 500 bp between pairs of start and end coordinates was allowed to take inaccurate breakpoint inferences into account. Furthermore, we normalized duplications to insertions since not all variant callers identify the same types of variants.

By default, Sniffles requires 10 supporting reads to call a structural variant. We also tested alternative minimum numbers of supporting reads to see the effect on accuracy. In addition, we performed a downsampling experiment of the alignment to see how it affects the performance of Sniffles. No parameter variation experiments were performed for NanoSV due to its long running times. We also evaluated the accuracy of the zygosity determination and investigated the difference in accuracy between "gain" and "loss" CNVs. All scripts for evaluation and plotting of our results are available on https://github.com/wdecoster/nano-snakemake/

### Structural variant analysis workflow

We generated a workflow for structural variant analysis based on the snakemake engine (Koster and Rahmann 2012), combining minimap2 (H.Li 2018) and ngmlr (Sedlazeck et al. 2018) for alignment followed by sorting and indexing bam files using samtools (H. Li et al. 2009) and structural variant calling using Sniffles (Sedlazeck et al. 2018) and NanoSV (Cretu Stancu et al. 2017). Per aligner, we took the union of the SV calls from Sniffles and NanoSV to form a high sensitivity set, and the intersection of both callers to form a high confidence set. Resulting variant files are processed using vcftools and bcftools (H.Li 2011; Danecek et al. 2011), combined using SURVIVOR (Jeffares et al. 2017) and annotated with information of segmental duplications, overlapping genes and known variants in the Database of Genomic Variants (DGV) (MacDonald et al. 2014) using vcfanno (Pedersen, Layer, and Quinlan 2016). Read depth is calculated using Mosdepth (Pedersen and Quinlan 2018). Plots were generated using Python scripts with the modules matplotlib (Hunter 2007), pandas (McKinney 2011), seaborn (Waskom et al. 2017) and cyvcf2 (Pedersen and Quinlan 2017). The snakemake workflow is available on https://github.com/wdecoster/nano-snakemake/. A graphical representation of the workflow can be found in Supplementary figure S1.

## Acknowledgments

The authors thank Jonathan Pugh and James Platt from Oxford Nanopore Technologies for support and troubleshooting when getting started with sequencing on PromethION, Jose Espejo Valle-Inclan for assistance in using NanoSV and interpreting its output, Fritz Sedlazeck for assistance in using Sniffles and SURVIVOR and Mark Chaisson for releasing the SV calls of NA19240. The study was in part funded by the VIB, a Life Sciences Institute in Belgium, the University of Antwerp, the Flanders Agency for Innovation and Entrepreneurship (VLAIO) and the VIB Tech Watch Fund. W.D.C. is a recipient of a PhD fellowship from IWT/VLAIO, A.D.R. is a recipient of a PhD fellowship from FWO.

## Disclosure declaration

W.D.C. and A.D.R. have received travel reimbursement from Oxford Nanopore Technologies to speak at London Calling 2018 and the ASHG 2018 meetings. Oxford Nanopore Technologies provided consumables free of charge for the realization of this project.

## Footnotes

1 ftp://ftp.ncbi.nlm.nih.gov/genomes/all/GCA/000/001/405/GCA_000001405.15_GRCh38/seqs_for_alignment_pipelines.ucsc_ids/GCA_000001405.15_GRCh38_no_alt_analysis_set.fna.gz

